# Effect of Lauric Acid on the Stability of A*β*_42_-Oligomers

**DOI:** 10.1101/2020.12.21.423843

**Authors:** Prabir Khatua, Asis Jana, Ulrich H. E. Hansmann

## Abstract

While Alzheimer’s disease is correlated with the presence of A*β* fibrils in patient brains, the more likely agents are their precursors, soluble oligomers that may form pores or otherwise distort cell membranes. Using all-atom molecular dynamics simulation we study how presence of fatty acids such as lauric acid changes the stability of pore-forming oligomers built from three-stranded A*β*_42_ chains. Such a change would alter the distribution of amyloids in the fatty-acid rich brain environment, and therefore could explain the lower polymorphism observed in A*β*-fibrils derived from brains of patients with Alzheimer’s disease. We find that lauric acid stabilizes both ring-like and barrel-shaped models, with the effect being stronger for barrel-like models than for ring-like oligomers.

## 1 INTRODUCTION

Long-term disturbance of brain function as seen in Alzheimer’s, Parkinson’s disease, or after traumatic brain injury, is often associated with the presence of amyloid fibrils in the brain; with the symptoms most likely caused by their precursors, small transient soluble oligomers formed by self-association of mis-folded proteins. In the case of Alzheimer’s Disease the amyloid deposits are rich in A*β* peptides, with the more frequent A*β*_40_ peptides observed in U-shaped configurations while A*β*_42_ can also take a three-stranded S-shaped motif. We have shown in earlier work^1,2^ that this geometry allows the more toxic A*β*_42_ to associate into assemblies which A*β*_1*−*40_ peptides cannot form. These include ring-like and barrel-shaped oligomers. The pore-like structure of both assemblies suggests water leakage through cell membranes as a possible mechanism for the higher toxicity of A*β*_42_. Formation of the oligomers and their propagation into mature fibrils are likely modulated by the brain environment as *in vitro* fibrils exhibit a rich polymorphism that is not seen in fibrils extracted from brain tissue of Alzheimer patients. ^3–5^ In Ref.^6^ we could indeed demonstrate that addition of lauric acid raises fibril stability for A*β*_42_, while no such stabilizing effect was found for A*β*_40_ fibrils. Our results further suggested that presence of lauric acid enhances the elongation of A*β*_42_-fibrils but not their nucleation. This is surprising as it has been shown that lipids and fatty acids can enhance oligomer formation. ^7–12^ Hence, one would expect also an effect on fibril nucleation, assuming that fibril growth starts with the size of the oligomers crossing a nucleation threshold.

Unfortunately, there is no easy way to probe in computational studies how fatty acids modify the formation and stability of the presumably more toxic oligomers. This is because these soluble oligomers exist in an ensemble of diverse and transient assemblies that are difficult to characterize by NMR or similar techniques, making it a challenge to obtain the structural models needed for computational stability investigations. An exception is the class of prion-like self-propagating large fatty acid-derived oligomers (LFAOs), extensively studied by the Rangachari lab.^13–15^ These assemblies of A*β*_42_ form in the presence of lauric acid but are stable even after removal of the fatty acids. It has been shown that in mice LFAOs cause cerebral amyloid angiopathy (also a common co-pathology in Alzheimer’s disease patients). ^16^ We have proposed in Ref.^17^ a ring-like structure as a model for the LFAOs that is indeed stabilized by presence of lauric acid. However, the effect was not as strong as needed to explain the experimental results of the Rangachari group. This result may either point to shortcomings in our model, which while consistent with experimental measurements of the Rangachari Lab^17^ may lack crucial details, or it may suggest that fatty acids shift the equilibrium toward toxic oligomers not by enhancing their stability but by a different mechanism. In order to probe this question, we extend in the present paper our earlier work^6^ and study in more detail the effect of lauric acid on A*β*_42_ oligomer models. Both ring-like and barrel-shaped models are considered as their pore-forming ability provides a potential mechanism for the presumed neurotoxicity of the oligomers. Going beyond the earlier work we include new a ring-like model where the individual chains take a slightly different and potentially more stable fold than considered previously, namely the one observed in the more recent Cryo-EM model of A*β*_42_. We also consider models with an out-of-register arrangement of the *β*_2_ and *β*_3_ strands, a motif that proved to be surprisingly stable in earlier work. ^2^ Our barrel-like models are similar to the cylindrin model proposed earlier as a model for toxic oligomers,^18^ however, unlike cylindrin the barrels not formed from six chains of a 13-residue peptides but built from either four or six A*β*_1*−*42_ chains. The stability of these models is studied in a series of molecular dynamics simulations, indicating that lauric acid indeed stabilizes the oligomers. The effect is more pronounced for barrel-like models than for ringlike oligomers.

## 2 MATERIALS AND METHODS

### 2.1 Model Construction

In a recent study^1^ we used an S-shaped model of A*β* peptides deposited in the Protein Data Bank under identifier 2MXU^19^ to construct N-fold ring-like assemblies, where two adjacent monomers in a layer are connected by an inter-chain salt bridge between residue K16 and residues E22 or D23, with the three *β*-strands arranged as in the in-register-fibril (IRF). In Ref.^17^ we proposed one of these ring-like assemblies, a six-fold double-layer (6×2) ring, as a structural model for the 12mers, observed by the Rangachari Lab as dominant species of their Large-Fatty-Acid-Catalyzed-Oligomers (LFAOs) in a low-concentration setting. Surprisingly, we did not find in Ref.^6^ a stabilization of this structure by the presence of lauric acid. For this reason, we have considered in the present study not only this model, but also the one where in the individual chains the *β*_2_ and *β*_3_ strands are in an out-of-register arrangement (ORF), replacing the inter-chain hydrogen bonds of the IRF geometry by intra-hydrogen bonds. The new models were derived by following the protocol described in.^2^ Note that unlike in our earlier work, all N-terminal residues have been added so that all our models are built from the full-sized A*β*_42_ chains and not only from fragments A*β*_1*−*42_. In addition we have also considered ring-like assemblies where the individual chains have the three-stranded *β*-sheet arrangement of the A*β*_42_ fibrils recently resolved by cryo-EM and deposited in the Protein Data Bank under identifier 5OQV.^20^ Both in-register and out-of-register arrangements of the *β*_2_ and *β*_3_ strands are considered. We refer to these arrangements as Model B, while the ones with the monomers in the S-shaped fold of the (PDB-ID:2MXU) fibril are called by us as Model A. The four models (Model A IRF and Model A ORF, Model B IRF and Model B ORF) were constructed by first using the VMD software^21^ to arrange in each case 12 chains into two layers with an approximate 60^*\**^-symmetry. Then we restrained the salt-bridge forming groups (COO^*−*^ and 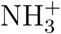) between neighboring chains using the NAMD code.^22^ Once the salt-bridges were formed, we released the restraints and continued the simulation for another ns. After a few such cycles, we arrived at the relaxed ring-like structures of Fig. 1a)-d) which we used as our start configurations in our stability simulations.

**Figure 1:**
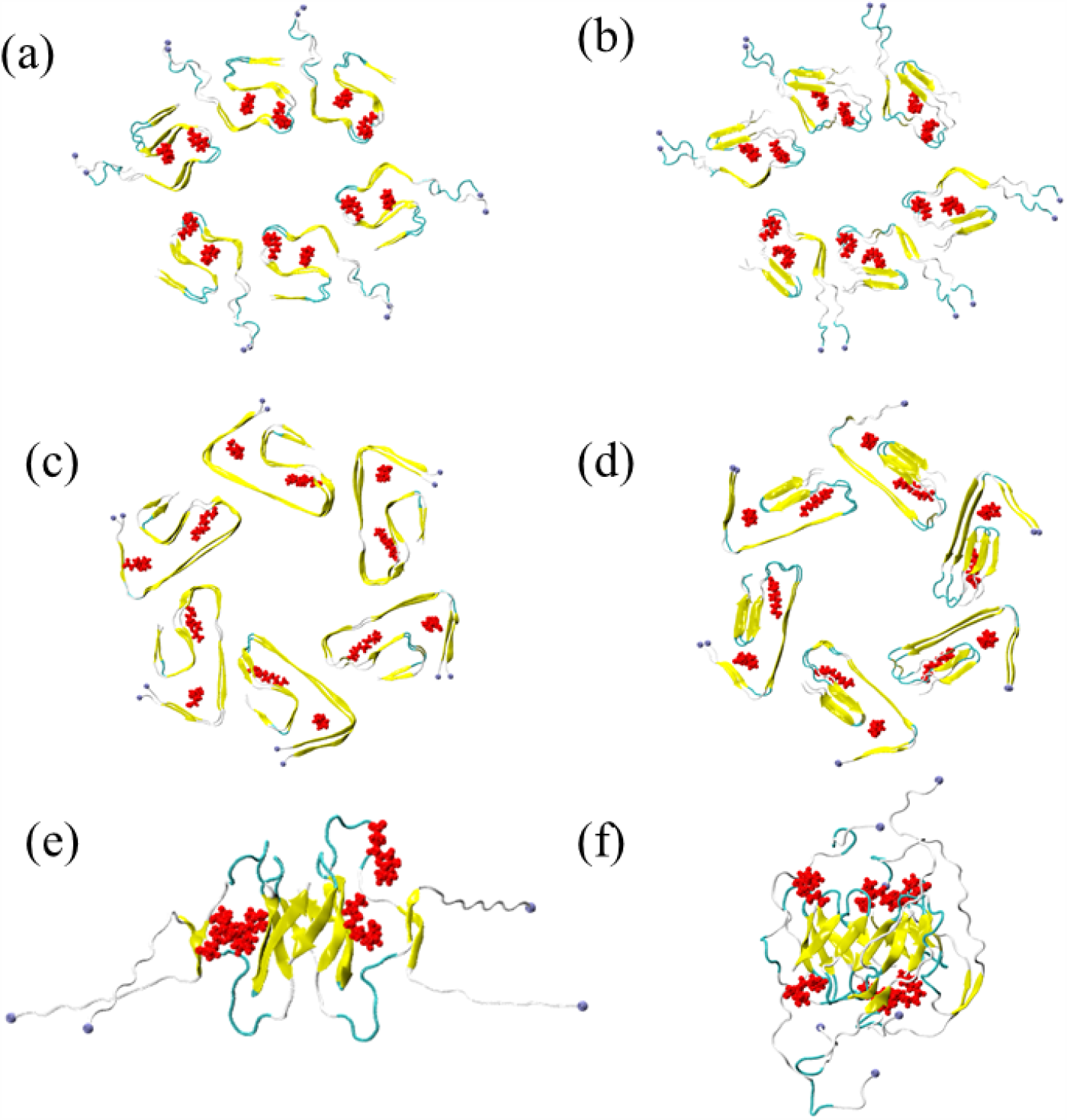
Start configurations for the ring-like model A oligomer with (a) in-register *β*_2_ − *β*_3_ strands and (b) out-of-register *β*_2_ − *β*_3_ strands; the ring-like model B oligomer with (c) inregister *β*_2_− *β*_3_ strands and (d) out-of-register *β*_2_ −*β*_3_ strands; (e) the barrel-shaped tetramer BB4 and (f) the barrel-shaped hexamer BB6. The systems are simulated both in absence and presence of lauric acid, with the binding sites of the fatty acids (as determined by Autodock) also shown in red color. The N-terminal ends of the individual chains are marked as spheres in blue.

In previous work,^2^ we could show that three-stranded A*β*_42_ chains with an ORF arrangement of the *β*_2_ and *β*_3_ strands can also form barrel-like oligomers. As the ring-like oligomer models discussed above, such barrels are again consistent with the pore-like A*β* assemblies observed by low-resolution atomic force microscopy. ^23^ We therefore consider also such assemblies in our stability study, focusing on the tetramer (BB4) and the hexamer (BB6), which were the smallest stable forms seen in our previous studies. For this purpose, we have used the same BB4 structure (built from N-terminal truncated chain fragments A*β*_11*−*42_) as prepared in our earlier study, ^2^ but adding to the individual chains the missing N-terminal residues 1-10. The resulting oligomer is shown in Fig. 1e). In a similar way, we construct the hexamer barrel by first placing six A*β*_27*−*42_ ORF fragments in such a way that neighboring chains are arranged in an anti-parallel fashion. This arrangement is one of an “unrolled” barrel and has the chains placed in such a way that the backbone atoms forming hydrogen bonds in the final barrel configuration^2^ are in close proximity. This unrolled hexamer configuration is again simulated with NAMD^22^ restraining the backbone hydrogen bond forming atoms. Once the appropriate hydrogen bond pattern is established after a few ns of simulation, the hexamer is rolled into a barrel, and simulated for a few more ns with restraints on the hydrogen bond forming atoms. Once the completely rolled A*β*_27*−*42_ hexamer barrel is formed, we relax the structure by releasing the restraints in a nother short simulation of a few ns. The remaining residues 1–26 are added to each of the monomers of this A*β*_27*−*42_ hexamer barrel. The full-length A*β* hexamer structure is once again simulated by applying the restraints on hydrogen bond forming residues in the core *β* strand region i.e., within *β*_2_ and *β*_3_ region for 1 ns followed by release of the restraints for another 1 ns. After a few cycles of such a simulation, we select the best BB6 configuration based on visual inspection and qualitative analysis of contact pattern of the structure. This selected configuration is used for further simulation of BB6 and doing docking, and is shown in Fig. 1f).

We have studied each of the above constructed systems both in presence and absence of lauric acid. When present, the ratio of lauric acid molecules and chain segments is 1:1, i.e., there are twelve lauric acid molecules added in case of ring-like models, four for the BB4 barrel, and six for the BB6 barrel. The lauric acid molecules were docked to the individual chains at sites determined by using the AutoDock software^24^ taking into account the docking score, visual inspection, and our previous knowledge of the systems. Assuming symmetric arrangements for the other chains, we then replicate the binding site for each monomer. Note that the purpose of the present investigation is to probe the stabilizing effects of lauric acid on the oligomer structure, not to identify the exact binding site. For this reason, we did not try to identify an optimal binding site but rather assume that possible better binding site will be found when starting from our guess. However, we nevertheless show the start binding sites also in Fig. 1 in the respective oligomer start configurations.

### 2.2 Simulation Set-up

Each of the systems is solvated in a cubic box of well-equilibrated water. The salt (NaCl) concentration is set to 0.1 M along with charge neutralizing each of the systems by adding required number of additional Na^+^ ions as all the systems studied in this study are negatively charged. While solvating the system with water, we have left a minimum of 1.2 nm distance between the edge of the box and the protein system. The number of water molecules, box dimension for each of the systems are given in Table 1. After initial minimization using the steepest descent algorithm as implemented in GROMACS software,^25^ each of the systems is simulated for a very short period of time with positional restraints on the protein atoms. The resulting configurations serve then as start point for the respective production run.

**Table 1:** The system details for the ring-like and barrel A*β*-oligomer structures.

| type | system | presence of lauric acids | dimension (nm) | no of water molecules |
| --- | --- | --- | --- | --- |
| Ring | Model A (IRF) | Yes | 17.3 | 1,67,712 |
|  |  | No | 17.3 | 1,67,858 |
| Ring | Model A (ORF) | Yes | 17.6 | 1,76,186 |
|  |  | No | 17.6 | 1,76,320 |
| Ring | Model B (IRF) | Yes | 15.3 | 1,14,793 |
|  |  | No | 15.3 | 1,14,966 |
| Ring | Model B (ORF) | Yes | 15.4 | 1,16,839 |
|  |  | No | 15.4 | 1,16,944 |
| Barrel | BB4 | Yes | 14.1 | 90,116 |
|  |  | No | 14.1 | 90,167 |
| Barrel | BB6 | Yes | 10.5 | 36,294 |
|  |  | No | 10.5 | 36,368 |

Our molecular dynamics simulations use the GROMACS software package.^25^ Protein-protein interactions are modeled with the CHARMM 36m force field,^26^ while water molecules are modeled with TIP3P,^27^ a combination that is known to perform well in studies of amyloid assemblies.^28^ The bond lengths are constrained using the LINCS algorithm, ^29^ while the SETTLE algorithm^30^ is utilized to maintain water geometry. Each of the systems is simulated in an isothermal-isobaric (NPT) ensemble at a temperature set to 310 K by a v-rescale thermostat,^31^ and pressure set to 1 bar by a Parrinello-Rahman barostat.^32^ The cut-off for electrostatic and van der Waals (vdw) interactions is set to 1.0 nm. Periodic boundary conditions (PBC) are used and the particle mesh Ewald (PME) method^33^ is employed to calculate the long-range electrostatic interactions. Equations of motions are integrated with a time step of 2 fs using the leapfrog algorithm as implemented in GROMACS.^25^ Each of the systems is simulated in triplicate (three independent runs for each systems). Each oligomer model is simulated both in presence and absence of lauric acids, with the later allowing to compare the change in stability. For the ring-like systems, the total simulation run length is 25 ns, while that for barrel systems is 200 ns. Configurations are saved every 50 ps along the trajectories.

### 2.3 Observables

The structural deviation is monitored by calculating the root-mean-square-deviation (RMSD) to the start configuration, while the residue-wise structural flexibilities are measured by calculating root-mean-square-fluctuation (RMSF).

The number of intra- and inter-monomer (side-wise and inter-layer in case of ring-like structures) contacts is calculated to probe the stability of the corresponding structure. Here the contacts are defined using a distance cut-off of 0.45 nm. Correlations between contacts are monitored by calculating the intermittent time correlation function (TCF), *C*_*contact*_(*t*), which is defined as^34–36^

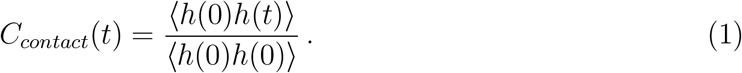

Here, *h*(*t*) is a population variable that takes a value of one if there exists a contact of the considered pair at a particular time *t*, and zero otherwise.

All our analyses are done by using our in-house codes, while the oligomer configurations are visualized using VMD software.^21^

## 3 RESULTS AND DISCUSSION

The underlying assumption of the present work is that the toxicity of A*β*_42_ aggregates is higher than that of the more common A*β*_40_ because A*β*_42_ peptides can form three-stranded motifs which in turn allow for assembly into pore-forming aggregates (ring-like or barrel-shaped); and that formation of these aggregates is enhanced by the brain-environment, specifically, the presence of fatty acids. In previous work we could show that presence of lauric acid stabilizes fibrils built from S-shaped A*β*_42_ chains but not fibrils formed by the U-shaped A*β*_40_ peptides. However, the disease-symptoms causing agents are likely not fibrils but soluble oligomers formed either on or off-pathway to fibril formation. There is some evidence that the neurotoxicity of the oligomers is related to their ability to form pores in the cell membrane.^37,38^ Unfortunately, these oligomers are transient and likely exist in a plethora of sizes, making it difficult to derive structural information by solid-state NMR, cryoEM or similar techniques. The Rangachari Lab could show that lauric acid catalyzes an ensemble of stable and homogeneous oligomers of either 12mers or 24-mers, depending on concentration and pH.^14^ They also could show that these self-propagating large fatty acid-derived oligomers (LFAOs) cause in mice cerebral amyloid angiopathy, a common co-pathology in Alzheimer’s disease patients. We have put forward in previous work^17^ a ring-like model for the 12mer LFAOs that is consistent with low-resolution AFR measurements taken in the Rangachari Lab. However, for us surprisingly, we did not find a noticeable stabilization of this assembly by lauric acid. For this reason, we extend here our earlier studies, considering now both ring-like and barrel-shaped as models for neurotoxic oligomers.

We begin our analysis by inspecting the ring-like oligomers proposed by us as models for the LFAOs studied in the Rangachari Lab. Visual inspection reveals that in absence of lauric acid both ring-like models, A and B, irrespective of whether the *β*_2_ and *β*_3_ strands are arranged in in-register-fibril (IRF) or out-of-register-fibril (ORF) decay very quickly (within 10 ns). This is despite the tight packing of the chains in the start configurations, with the maximum number of possible inter-layer hydrogen bonds holding the two layers together, and K16-E22/D23 salt-bridges connecting adjacent chains in each layer. The fast decay despite the presence of multiple and strong contacts suggests that the short life-times of the two-layered *β*-sheet arrangements are due to inherent chain-flexibility, leading to the destruction of the packing. If this assumption is correct, then the experimentally observed catalysis of A*β*_42_ chains into the LFAOs implies a mechanism by which the flexibility of the chains is reduced.

In order to check this hypothesis we have also simulated the four ring-like models in the presence of lauric acid. While in all cases the complex of A*β*_42_ ring-like oligomers with lauric acid decays again on a very short timescale, we do find now a stabilizing effect by lauric acid. For instance, in Figure 2 we show the time evolution of the number of native inter-layer and side-wise inter-chain contacts for model A, and compare the data between the two cases i.e., in presence and absence of lauric acids. In order to ease comparison, we have normalized the number of contacts in the start configurations to one. In the cases where the *β*_2_ and *β*_3_ strand are in-register and stabilized by inter-layer hydrogen bonds, see Fig. 2a)-b), both the number of native inter-layer and side-wise contacts stays higher when lauric acid is present and binding to the rings. Note, however, that both in presence and absence of lauric acid the number of both contacts decreases rapidly within the first ns. This decrease is due to break-up of the rings, which happens at random places depending on the trajectory (i.e., the strain on the structure is not concentrated at a certain position), but here in both layer at the same time. The loss of side-wise contacts at the break-up position is also correlated with a loss of inter-layer contacts in the neighborhood of the break-up point. Once, the ring breaks up at a given point, it deteriorates further, however, now slower in presence of lauric acids than in absence. The effect is more pronounced for the side-wise contacts than for the inter-layer contacts, suggesting that while binding of lauric acid does not prevent the initial breakup of the ring geometry, it encourages packing of chains over their stacking. However, the contact-forming role of lauric acid is non-specific, and it also encourages formation of non-native contacts along the trajectory. This non-specific effect is also seen when in model A the *β*_2_ and *β*_3_ strands are out of register, with the hydrogen-bonding now intra-chain, see Fig. 2c)-d. On the other hand, no stabilizing effect is seen when considering only the native contacts.

**Figure 2:**
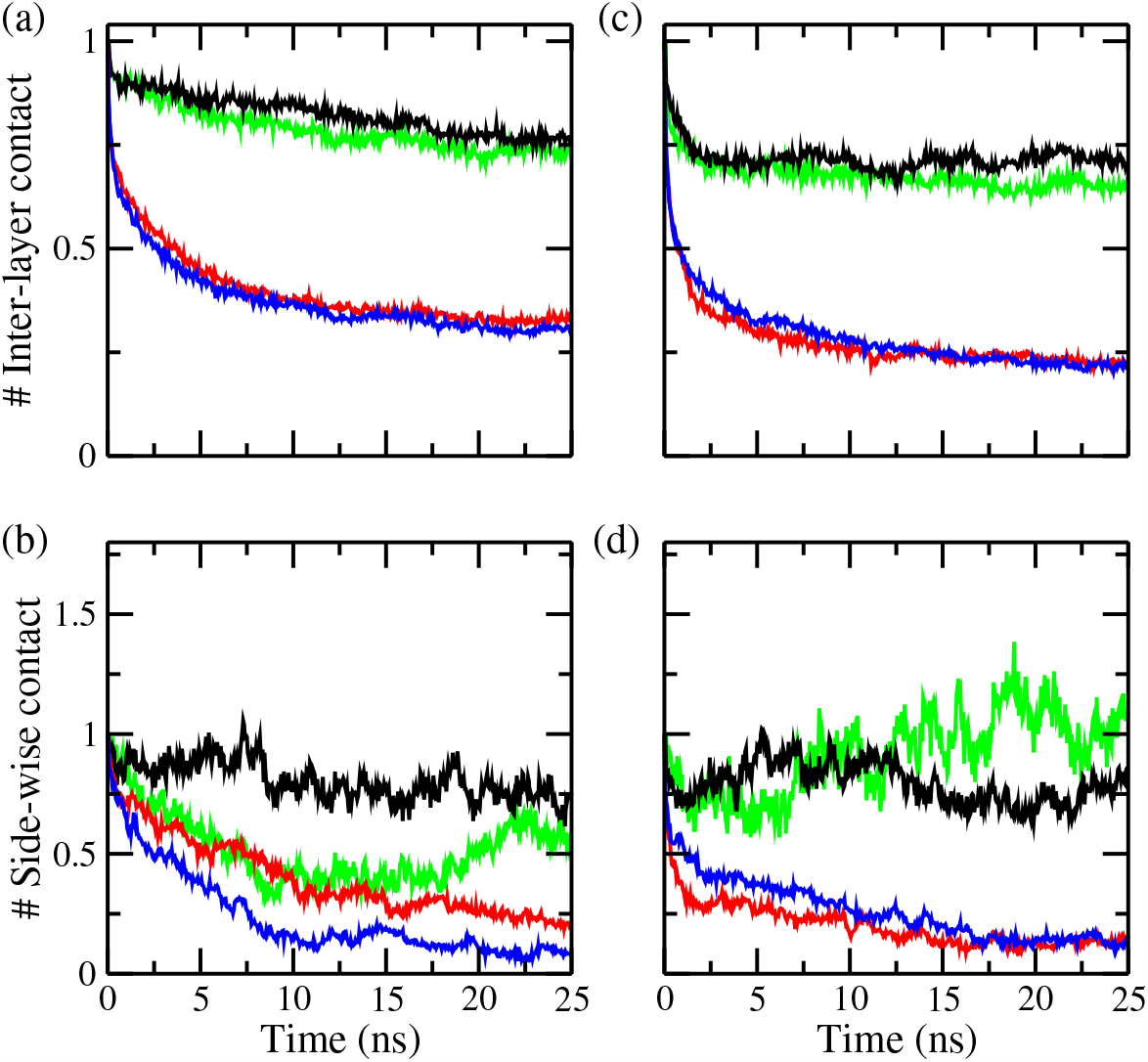
Average number of inter-layer and side-wise (within a layer) inter-chain contacts as a function of time for Model A ring-like oligomers with (a-b) in-register-fibril (IRF) or (c-d) out-of-register-fibril (ORF) *β*_2_ − *β*_3_-sheet arrangements. Averages are taken over all three runs simulated for each system, and normalized in such a way that the corresponding value at start is the unity. The results for the *total* number of contacts are shown in black (in presence of lauric acids) and green (in absence of lauric acids), while the corresponding numbers of *only native* contacts are displayed in red (in presence of lauric acids) and blue (in absence of lauric acid).

Similar to what was seen by us in earlier work, ^6^ the ring-like oligomers of Model A type decay even in the presence of lauric acid and for in-register arrangement of the *β*_2_ and *β*_3_ strands much faster than the experiments in the Rangachari lab would let one expect. The situation is worse for Model B, where the individual chains take a form as seen in the Cryo-EM model (PDB ID: 5OQV)^20^ of an A*β*_42_ fibril. Data are shown in Fig. 3. For the in-register model, no clear signal is seen for stabilization of side-wise contacts, and only a weak one for the inter-layer contacts. Again, no significant difference is seen in the out-of-register case systems with lauric acid present and such without lauric acid. Unlike to model A we do not see an effect of lauric acid on the formation of non-native contacts. Comparing the two models, we note that the initial two binding positions as obtained from docking are not the same in the two models. In model A, the first binding position is in the region spanned by L17 to N27, while the second one is at A30–I32. On the other hand, the first binding site for model B is within the cavity formed by F4–H14 and L17–F19, and the second one is located within N27–A30. This difference in binding sites may explain the weaker stabilization by lauric acid.

**Figure 3:**
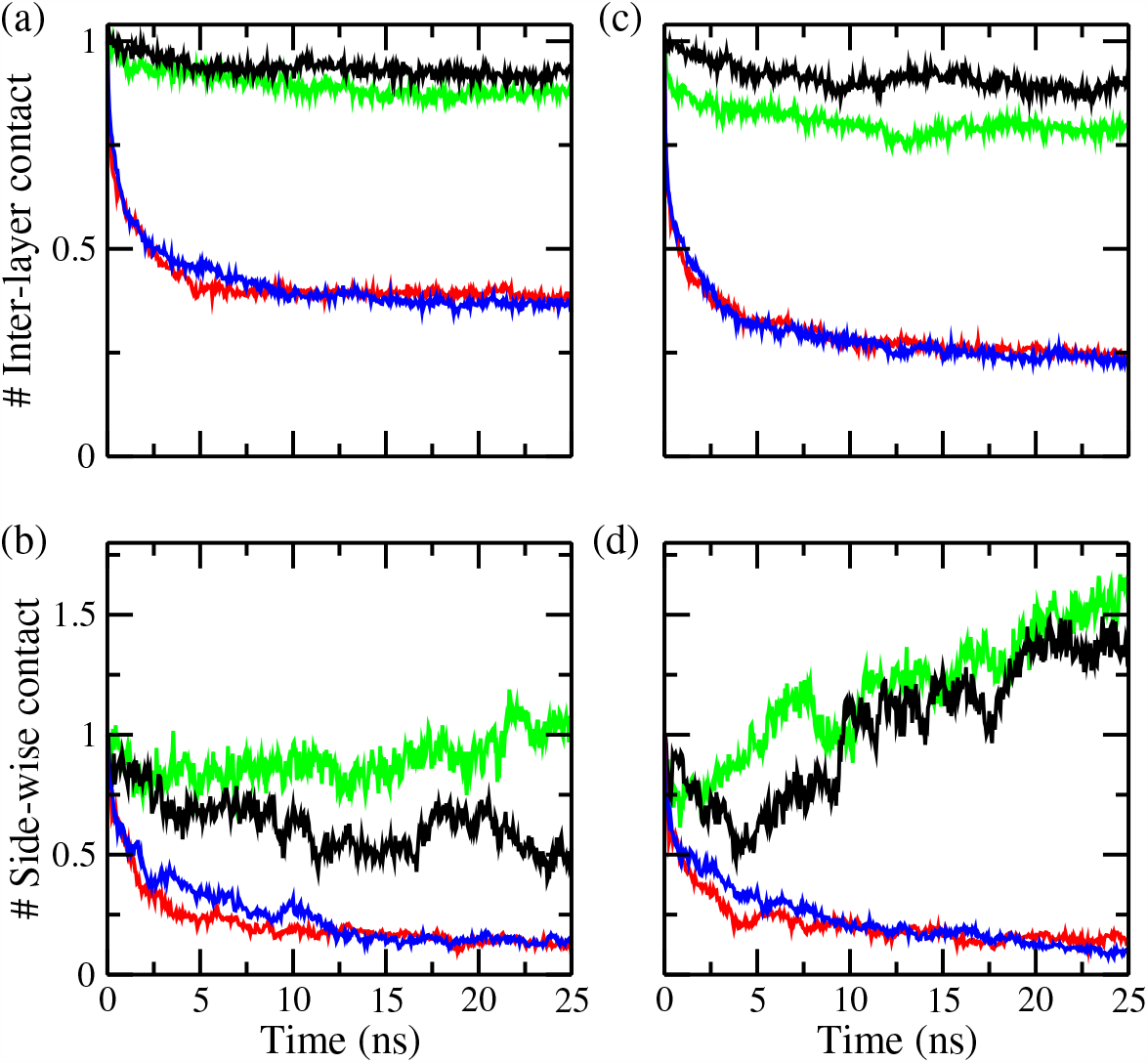
Average number of inter-layer and side-wise (within a layer) inter-chain contacts as a function of time for model B ring-like oligomers with (a-b) in-register-fibril (IRF) or (c-d) out-of-register-fibril (ORF) *β*_2_− *β*_3_-sheet arrangements. Averages are taken over the three runs simulated for each system, and normalized in such a way that the corresponding value at start is the unity. The results for the *total* number of contacts are shown in black (in presence of lauric acids) and green (in absence of lauric acids), while the corresponding numbers of *only native* contacts are displayed in red (in presence of lauric acids) and blue (in absence of lauric acid).

Note that in the case of Model A, the stabilization of side-wise contacts is more pronounced than that of the inter-layer contacts. This is likely because the stacking is already stabilized strongly by the inter-layer hydrogen bonds, making any stabilizing effect more difficult to detect than for the side-wise contacts. Reducing the inter-layer contacts by switching from in-register arrangement of the *β*_2_ − *β*_3_ strands with inter-layer hydrogen bonding to the out-of-register arrangement, where the hydrogen bonding is now intra-chain, increases the flexibility of the chains, and therefore reduce further the stability of the rings, likely to an extent that cannot be overcome by the stabilizing effect of lauric acid. Note that the lauric acid molecules do not stay at their initial binding sites after break-up of the rings. Instead, they diffuse along the surface, or even move transiently away from the surface, often returning at a different site. This movement of the lauric acid molecules may explain the loss of native contacts (at the original binding sites) and formation of new non-native contacts (at the new binding sites).

While the stabilizing effects of lauric acid on the ring-like assemblies cannot explain formation and long lifetimes of the LFAOs, they confirm that lauric acid can stabilize oligomers, and therefore may play a role in the self-assembly of A*β*_42_ aggregates. As our results raise doubts on ring-like structures as the motif for toxic A*β*_42_ oligomers we have looked into an alternative motif, namely barrel-shaped assemblies. Here, we have studied the tetramer (BB4) and hexamer (BB6) A*β* barrel which we found in earlier work to be the smallest stable barrel-shaped assemblies. Our goal is to investigate how lauric acids stabilize the assembly and how it encourages the self-assembly of these oligomers. Figure 4 depicts the root-mean-square-deviation as function of time, calculated over all backbone atoms of all chains and evaluated with respect to the corresponding start configuration. An increase in stability in the presence of lauric acids is clearly visible as the RMSD values of the systems with lauric acid are lower than found in the corresponding control systems (the ones without lauric acid). This is particularly true for BB6. The results also show that RMSD values approach a plateau after 100 ns. Therefore, we discard the first 100 ns and use the rest of the 100 ns trajectories for further analysis.

**Figure 4:**
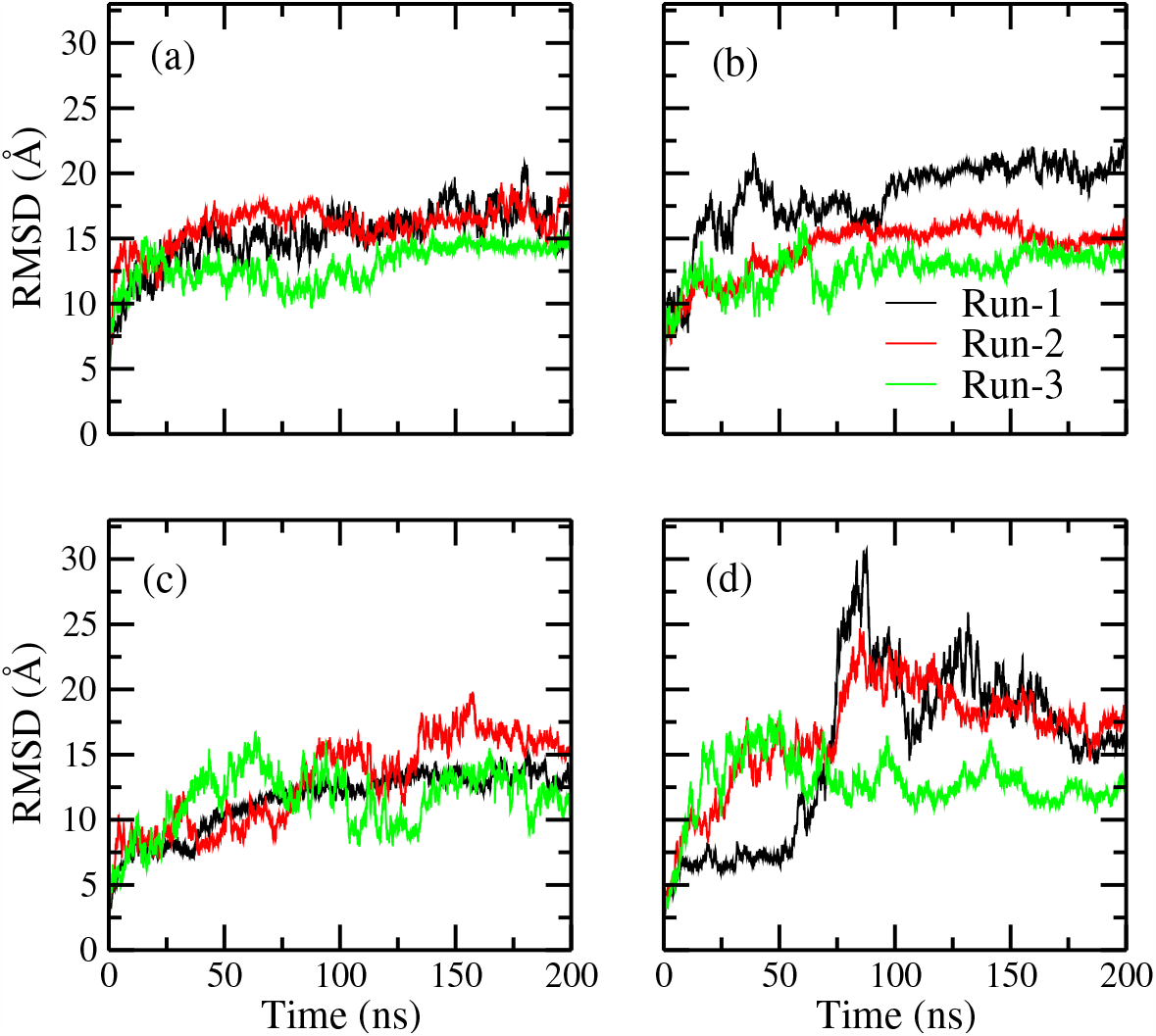
Root-mean-square-deviation (RMSD) to the start configuration as a function of time for the tetramer barrel (BB4) (a) in presence of lauric acid and (b) in absence of lauric acid. Results for the hexamer barrel (BB6) in presence of lauric acid are shown in (c), and the ones taken in absence of lauric acid are shown in (d).

The differences in absolute RMSD values and how they approach the equilibrium suggest that presence of lauric acid alters the structure of the barrel configurations. In order to check these structural changes, we have visually inspected configurations along the trajectories. Snapshots of these configurations as obtained at the end of one of the simulations are presented in Figure 5 for each case. Comparing Figure 5a) and b), one finds only little differences between BB4 configurations in presence and absence of lauric acids. This visual observation consistent with the fact why there was not much difference in the RMSD values between the cases. BB4 configurations are stable irrespective of the presence of lauric acids. However, the structures are found to be a little bit more distorted in absence of lauric acids. Presence of lauric acid helps reduce this distortion by facilitating the rearrangement of the A*β*-monomers to have an optimum packing and consequently to form a more optimal barrel. The role of lauric acids in stabilizing the barrel is clearly noticeable for larger barrels, as the hexamer barrel in Figure 5d) is more stable than the one in Figure 5e) for the system without lauric acid which shows a tendency to decay into monomers.

**Figure 5:**
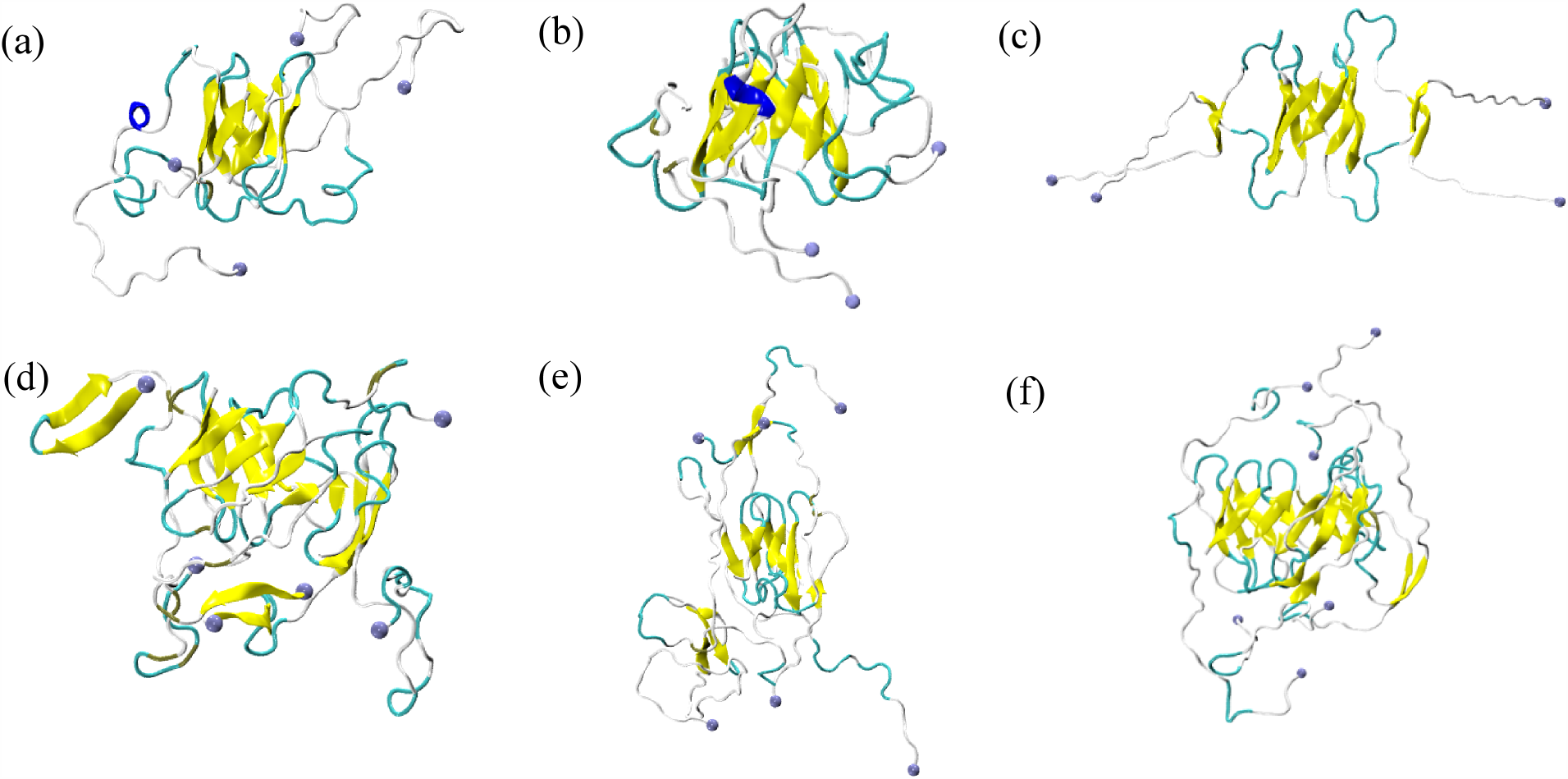
Representative snapshots as obtained at the end of a 200 ns simulated trajectories for the tetramer barrel (BB4) (a) in presence or (b) absence of lauric acids. For comparison, we show also the start configuration in (c). Similar configurations are shown for the hexamer (BB6) in (d) -(f), respectively. The N-terminals of each of the individual chains are marked as blue spheres.

Next, we have investigated how individual residues of the barrel get influenced by the presence of lauric acids, looking at the residue-wise root-mean-square-fluctuation (RMSF) as shown in Figure 6. This quantity is a measure for the flexibility of the system at the location of the specific residue, and exhibits a clear signal for the presence of lauric acid. The signal further differs between tetramer and hexamer. For the larger BB6 hexamer the presence of lauric acid leads to a decrease in the structural fluctuations, consistent with the stabilizing effect of lauric acids observed in the RMSD evolution. On the other hand, presence of lauric acids results in higher structural fluctuation in the case of BB4. This apparently surprising result can be explained by the fact that presence of lauric acids encourages the individual residues of each chain to rearrange as to form a more optimized barrel (see Figure 5), i.e., correcting a less-than-optimal geometry of the start configuration. This structural rearrangement of the residues is responsible for the higher RMSF values observed in the simulations that included lauric acid when compared to simulations without lauric acid added. Therefore, while we notice a stabilizing effect of lauric acid, the way these fatty acids stabilize the barrel geometry for BB4 and BB6 differs: for the smaller barrel (BB4), which is already stable due to proper packing, lauric acid facilitate monomer rearrangement so as to optimize the barrel geometry; for the larger barrel (BB6), which has a tendency to decay into monomers, lauric acid lowers the risk of this decay by reducing the structural fluctuations of the individual residues.

**Figure 6:**
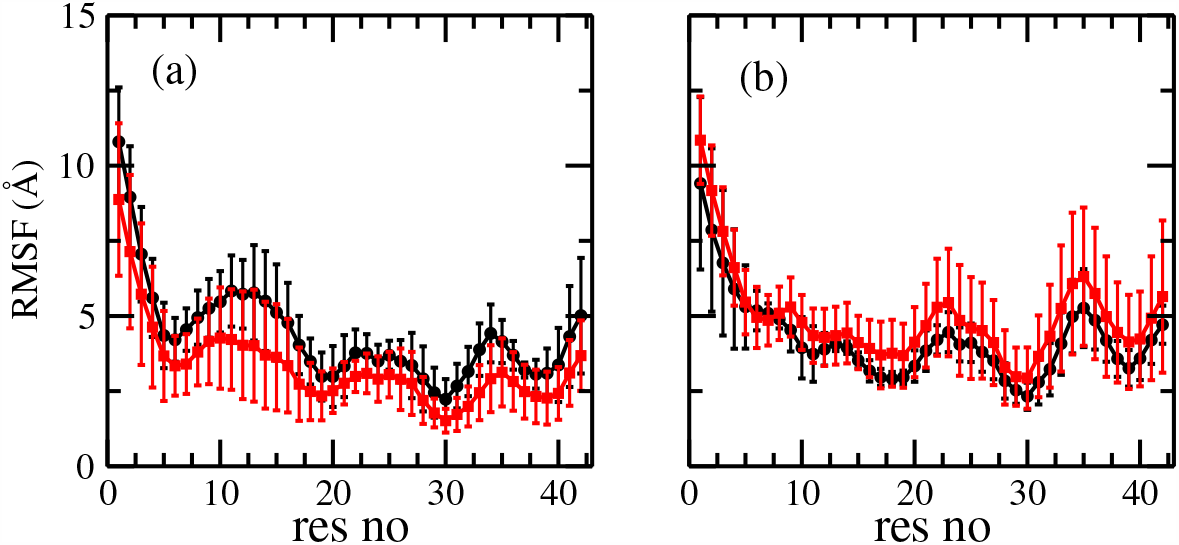
Residue-wise root-mean-square-fluctuations (RMSF) of (a) tetramer (BB4) (b) hexamer (BB6) barrel structures formed by A*β*_42_ peptides in the presence (black) or in the absence (red) of lauric acid molecules bound to the start configuration. The vertical lines show the error bar as calculated over the three independent simulations.

We have discussed so far how lauric acid stabilizes the barrels by by either restricting decay into monomers or by optimizing the barrel geometry. However, the results presented above are qualitative. In order to measure quantitatively the structural distortion of the barrel geometry, one needs to define a parameter that can quantify the distortion. For this purpose, we decided to measure the pore diameter of the barrels. Since we are interested in the barrel geometry, we only focus on the core region which is formed by residues spanning from N27 to A42. Here, the center-of-mass distance between two oppositely facing A*β* monomers is used by us to define the barrel diameter. This leads to two (between monomer pairs 1-3, *D*_13_, and 2-4, *D*_24_) or three (between monomer pairs 1-4, *D*_14_, 2-5, *D*_25_, and 3-6, *D*_36_) such pore diameters for BB4 and BB6, respectively. Hence, we define the extension of a pore by the average of the two (three) distances, while the deviation from barrel-shape is captured through a pore distortion parameter *D*_*p*_ defined for the tetramer BB4 by

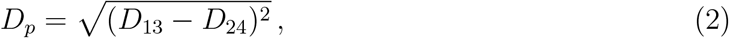

and for the hexamer BB6 by

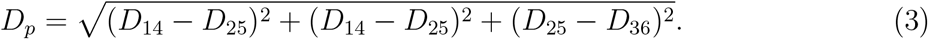

With this definition, we find as pore size for the tetramer BB4 values of 12.1 (0.5) Å in both cases (with and without lauric acid). The corresponding values for the hexamer are 19.7 (0.9) Å and 23.2 (7.4) Å, respectively, and show a larger difference between the barrel complexed with lauric acid, and the one without lauric acid. A similar but more pronounced behavior is found for the distortion of the pore. This can be seen in Figure 7 where we display the distribution of the deviation of the pore distortion parameter *D*_*p*_, as obtained from last 100 ns of trajectories and averaged over three independent runs. For comparison, the results as obtained in the absence of lauric acids are also shown in the figure. We remark that the bimodal nature of the distribution is an artifact of the small number of runs: in BB4 one of the trajectories for BB4 in complex with lauric acid got more strongly distorted than the other two runs, while the same happened for the hexamer BB6 in the case of absence of lauric acid. However, even without these outliers, the main picture stays unaltered: presence of lauric acid leads for the tetramer BB4 to a shift in the distortion parameter towards larger values, while the opposite is seen for the hexamer BB6. Note that the effect for BB4 is small, and visual inspection rather suggests a less distorted barrel geometry than seen in absence of lauric acids. Unlike BB4, BB6 configurations have more distorted pores as evident from the distribution of the distortion parameter *D*_*p*_. This is true irrespective of the presence of lauric acids, suggesting that BB6 configurations are not as symmetric as BB4 is. However, the shift of the curve towards the lower value once again signifies that lauric acids play a significant role in helping them retain the barrel geometry. In fact, in absence of lauric acids, BB6 configurations have a tendency to decay into monomers, as the curve shifts to a very high value especially for *D*_14_ and *D*_26_, indicating significant distortions of barrel geometry.

**Figure 7:**
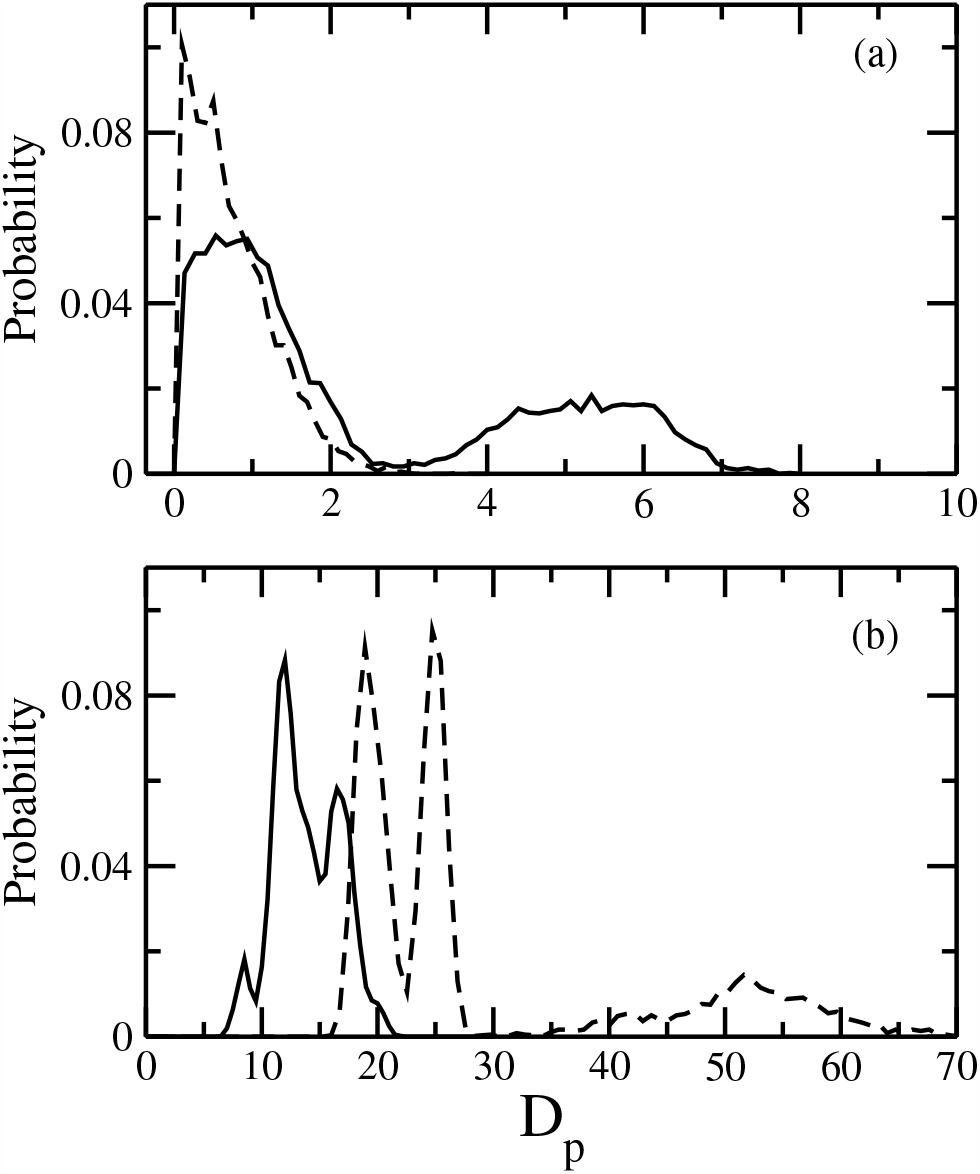
Distribution of pore distortion parameter (*D*_*p*_, as defined in the text) for (a) the tetramer (BB4) and (b) the hexamer (BB6) barrel as obtained from the last 100 ns trajectories of all three trial runs. The solid line represents data in the presence of lauric acids, while that with dashed line indicates the data in the absence of lauric acids.

Although there is a stabilizing effect of lauric acids, we have observed contrasting mechanisms for BB4 and BB6. In case of BB4, individual chains undergo a conformational rearrangement in presence of lauric acids before eventually transforming into a less distorted barrel. On the other hand, presence of lauric acids restricts the decay of BB6 into monomers and thus help them retain the barrel geometry. In order to scrutinize this contrasting behavior of BB4 and BB6 in more detail, we have calculated the average number of intra- and inter-monomer side-chain contacts, considering either all residues (residue 1–42), only the segments made of residue 11–42 (the region with all three *β*-strands), or restricting us even further to the core region of the barrel by considering only the segments made of residues 27–42. The distribution of the contacts as obtained from the last 100 ns trajectories averaged over three runs is shown in Figure 8. Once again we also add for comparison the data as measured in the control simulations where no lauric acid was added.

**Figure 8:**
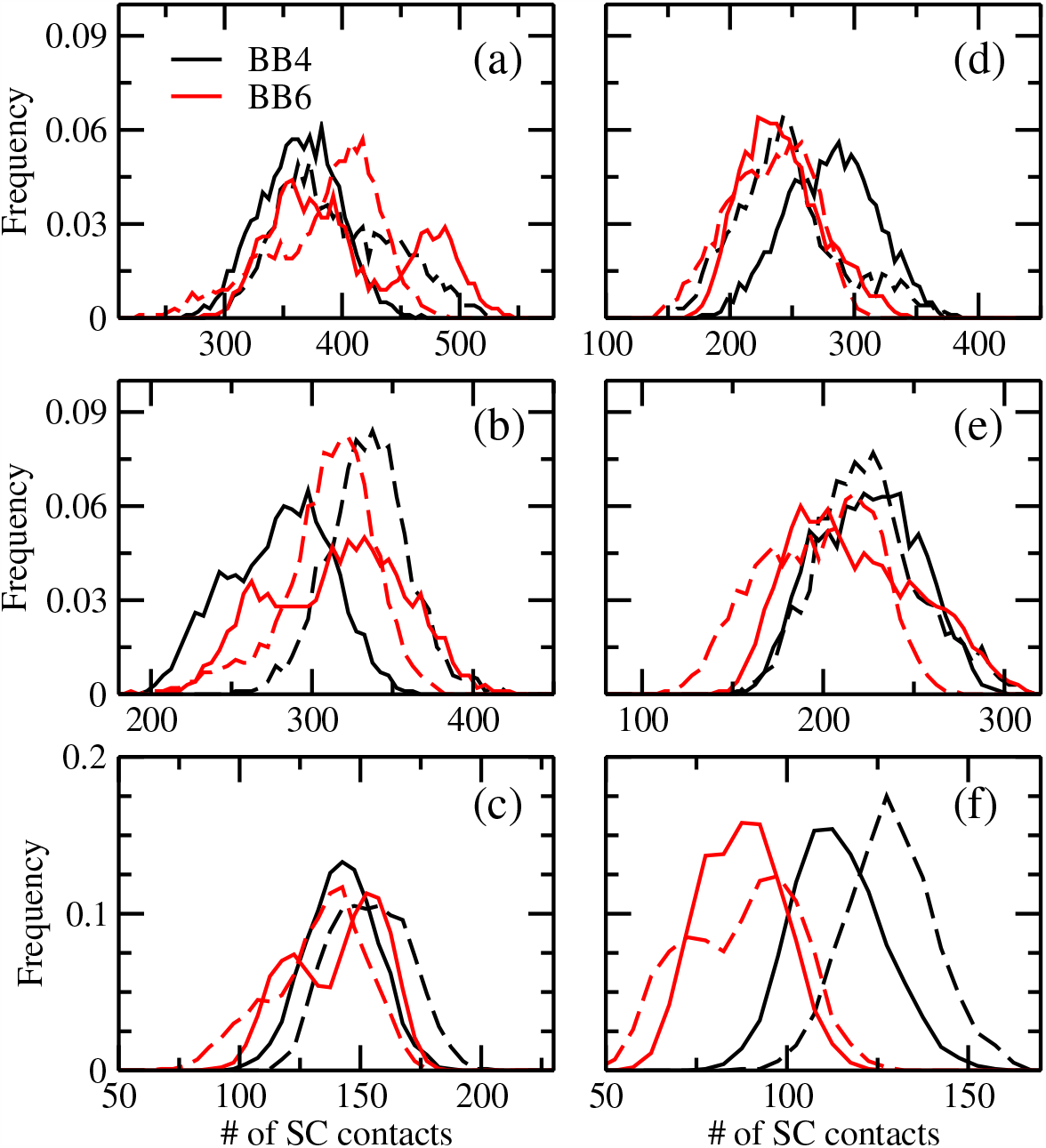
Distribution of number of (a-c) intra-monomer and (d-f) inter-monomer side-chain contacts for the tetramer BB4 (shown in black) and the hexamer BB6 (shown in red). The solid line represents the data in presence of lauric acid, while that with dashed line indicates the data in absence of lauric acids. The results are presented considering different parts of the monomers: top row (all residues), middle row (residues 11–42), and bottom row (residues 27–42)

As can be seen from the figure, both the number of intra- and inter-monomer side-chain contacts decrease in presence of lauric acids for BB4. The effect is most pronounced for the core region (residue 27-42). This decrease is correlated with the increase in flexibility of residues for BB4 (see also Figure 6(a)). On the other hand, on an average the number of both intra- and inter-monomer side-chain contacts increase in presence of lauric acids for BB6. While the increase in side-chain contacts in presence of lauric acids for BB6 can explain the observed stabilization of the BB6 geometry, the decrease in the same quantity in BB4 contradicts the idea that lauric acid stabilizes the BB4 barrel. We argue that presence of lauric acids disturbs the contact pattern in BB4 by forcing the individual monomers to rearrange into a structure with smaller number of contacts but more stronger interactions than seen in absence of lauric acids. This hypothesis is supported by the Figure 9, where we show the intermittent time correlation function for the side-chain contacts, *C*_*contact*_(*t*). The decay of intra-monomer side-chain contacts is faster in presence of lauric acids for BB4, suggesting shorter lifetime of those contacts, whereas the opposite is the case for inter-monomer sidechain contacts. Note that number of inter-monomer side-chain contacts is for BB4 lower in presence of lauric acids than that in absence of lauric acids (Figures 8(d-f)). This indicates that lauric acid weakens the strength of intra-monomer side-chain contacts in BB4, therefore enabling movements of the chains that lead to formation of stronger inter-monomer contacts. Thus, the overall result is a stabilization of the barrel geometry. On the other hand, in case of BB6, the decay is faster in absence of lauric acids irrespective of side-chain contact types, which simply explains why BB6 is more stable in presence of lauric acids.

**Figure 9:**
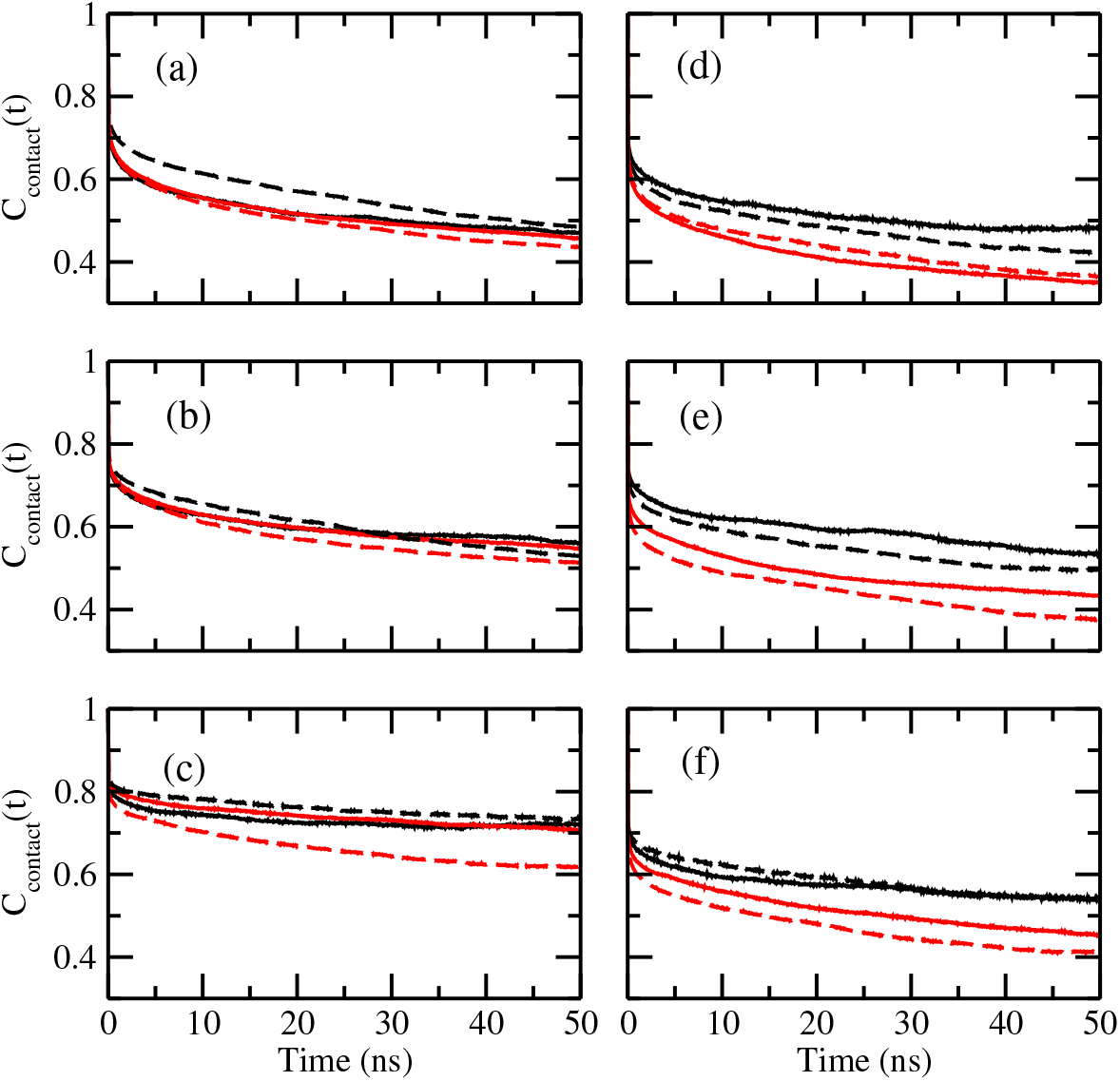
Contact correlation function for (a-c) intra- and (d-f) inter-monomer side-chain contacts. Data for the tetramer (BB4) are drawn in black, and such for the hexamer (BB6) in red. Data from systems simulated in the presence of lauric acid are drawn as solid lines, while such from simulations without added lauric acid are drawn as dashed lines. Results are shown considering either all residues (top row), or only residues 11-42 (middle row), or residues 27-42 (bottom row).

Finally, we investigate the binding of lauric acid to the barrels in order to understand better their lauric acid enabled stabilization. As the initial binding sites are obtained from docking, they may not be the optimal choices, and over the course of the simulation lauric acid may try to find a better binding pose. Indeed, visual inspection of configurations shows that in both BB4 and BB6 oligomers the ligands do not stay bound to a particular site; rather they move along the surface of each of the chains. In some cases they even leave the barrel surface, and when after some time returning, attach to a different chain and/or binding site. This movement is connected with a structural rearrangement of the chains leading to a stabilization of the barrel by increasing both lauric acid-barrel contacts and contacts within or among the A*β* chains. Note that the lauric acid molecules tend to return to the same residue (not necessary on the same chain) as they docked to in the start configuration. We have verified this visual impression tby calculating and comparing residue-wise binding probabilities as shown in Fig. 10. For this purpose, we define a binding site as the closest residue with at least one non-hydrogen atom within 0.45 nm from the lauric acid molecule. While we find for both BB4 and BB6 barrel for all residues non-zero probabilities (i.e., lauric acid molecules are seen transiently in the vicinity of all residues), there are two predominant binding sites: R5 and K28. These two residues (both charged and basic) are close to the initial binding site. Hence, the structural re-arrangement and movement of lauric acid molecules do not result from sub-optimal binding sites found by our docking algorithm.

**Figure 10:**
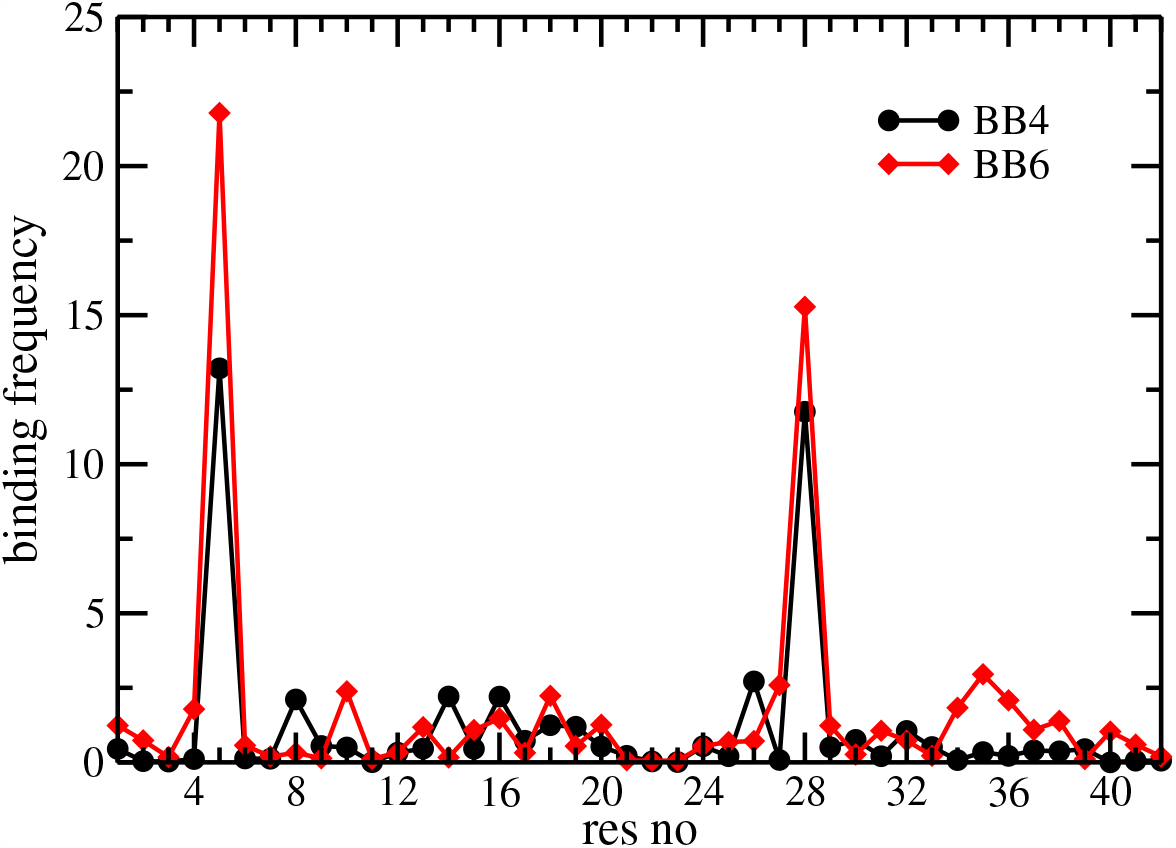
Residue-wise binding probability (normalized) of lauric acid for the tetramer (BB4) and hexamer (BB6) barrel structures. Data are calculated from the last 100 ns of all three independent runs in which the respective system was simulated.

Note that the binding probability is higher for BB6 than for BB4, indicating stronger binding of lauric acids to the hexamer. We conjecture that the role of lauric acids is to stabilize the barrel structure, and once a barrel is sufficiently stable, presence of lauric acid is no longer required. In this sense, lauric acid does catalyze the formation of the barrel. Hence, as the hexamer BB6 is less stable than the tetramer BB4, lauric acid molecules need to bind stronger and for longer time to stabilize the barrel geometry. On the other hand, the BB4 tetramer start configuration is already stable, and thus lauric acid just optimizes this motif.

## 4 CONCLUSIONS

Using molecular dynamics we have studied how presence of lauric acid alters the stability of pore-forming oligomers built from three-stranded A*β*_42_ peptides. Our investigation is motivated by the hypothesis that the brain environment, and here specifically the abundance of fatty acids such as lauric acid, alters the distribution of oligomers and fibrils, resulting in higher neurotoxicity and lower polymorphism than observed in vitro. As candidate for the neurotoxic oligomers we have simulated both ring-like and barrel-shaped assemblies. In all cases we find that presence of lauric acid stabilizes the oligomers, however, the effect is surprisingly small for our ring-like models. This may either point to shortcomings in our model, which while consistent with experimental measurements of the Rangachari Lab, ^17^ may lack crucial details, or it could suggest that fatty acids shift the equilibrium toward toxic oligomers by a different mechanism than enhancing the stability of the oligomers.

On the other hand, the stabilizing role of lauric acid is more pronounced for barrel-shaped assemblies. Here, lauric acid either stabilize an existing motif and inhibits decay into monomers (for the hexamer BB6), or causes structural re-arrangements that lead to more stable geometries (in case of the tetramer BB4). In both mechanisms is the position of the lauric acid molecules not static. Instead, the molecules transiently disconnect from the barrels only to return later to their binding sites (not necessary on the same chain). Binding of lauric acid to the start poses is higher for the hexamer BB6 than for the tetramer BB4, and as a consequence presence of lauric acid increases in the hexamer both number and life time of stabilizing contacts. We conjecture that lauric acid catalyzes formation of barrel-like assemblies by increasing their stability. Once a barrel is sufficiently stable, presence of lauric acid is no longer required to preserve its structural integrity. Hence, for the tetramer BB4, where the lauric acid molecules bind less strongly to their binding sites, we do not find an energetic stabilization. Instead, the transient movement of lauric acid rather leads to an increased flexibility of the residues in the A*β*_1*−*42_ chains that allow for rearrangements of structure and contact pattern which in turn optimizes the barrel and its stability. Taken together, our present study shows that presence of fatty acid can catalyze formation of certain A*β*-oligomers (in our case barrel shaped tetramers and hexamers built from of three-stranded A*β*_1*−*42_ chains) although the mechanism of motif stabilization may differ and depend upon the size of barrel.

## Acknowledgement

We thank Elliot K. Vanderford who helped us set up the systems with docking at the initial phase of this project. The simulations in this work were done using the SCHOONER cluster of the University of Oklahoma and XSEDE resources allocated under grant MCB160005 (National Science Foundation). We acknowledge financial support from the National Institutes of Health under grant GM120634 and AG062292.

## Graphical TOC Entry

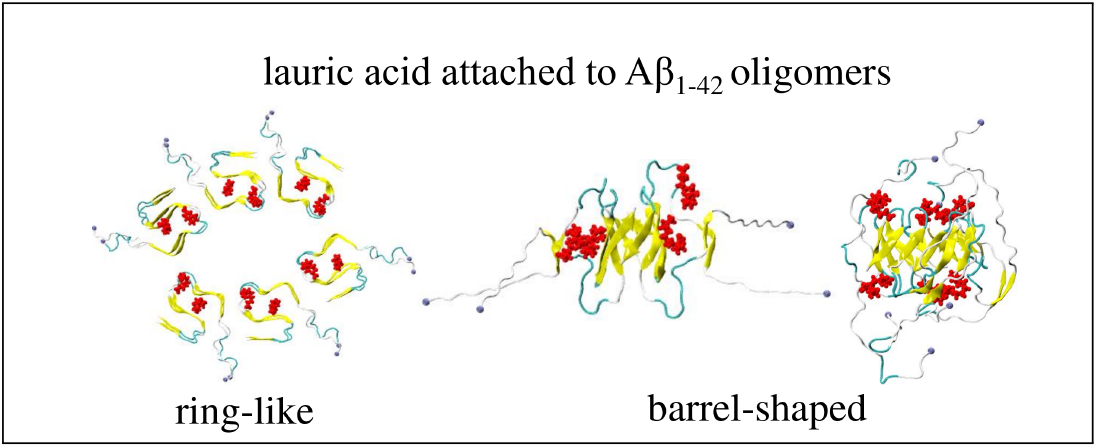

